# Inducible mucosa-like differentiation of head and neck cancer cells drives the epigenetically determined loss of cell malignancy

**DOI:** 10.1101/2023.06.30.547265

**Authors:** Felix Oppel, Sarah Gendreizig, Laura Martinez-Ruiz, Javier Florido, Alba López-Rodríguez, Harkiren Pabla, Lakshna Loganathan, Leonie Hose, Philipp Kühnel, Pascal Schmidt, Matthias Schürmann, Judith Martha Neumann, Flavian Viyof Ful, Lars Uwe Scholtz, Dina Ligum, Frank Brasch, Karsten Niehaus, Germaine Escames, Tobias Busche, Jörn Kalinowski, Peter Goon, Holger Sudhoff

## Abstract

**Background:** Human papillomavirus-negative head and neck squamous cell carcinoma (HNSCC) is a highly malignant disease with high death rates that have remained substantially unaltered for decades. Therefore, new treatment approaches are urgently needed. Human papillomavirus-negative tumors harbor areas of terminally differentiated tissue that are characterized by cornification. Dissecting this intrinsic ability of HNSCC cells to irreversibly differentiate into non-malignant cells may have striking tumor-targeting potential.

**Methods:** We modeled the cornification of HNSCC cells in a primary spheroid model and analyzed the mechanisms underlying differentiation by RNA-seq and ATAC-seq. Results were verified by immunofluorescence using human HNSCC tissue of distinct anatomical locations.

**Results:** HNSCC cell differentiation was accompanied by cell adhesion, proliferation stop, diminished tumor-initiating potential in immunodeficient mice, and activation of a wound healing-associated signaling program. Small promoter accessibility increased despite overall chromatin closure. Differentiating cells upregulated KRT17 and cornification markers. Although KRT17 represents a basal stem-cell marker in normal mucosa, we confirm KRT17 to represent an early differentiation marker in HNSCC tissue and dysplastic mucosa. Cornification was observed to frequently surround necrotic and immune-infiltrated areas in human tumors, indicating an involvement of pro-inflammatory stimuli. Indeed, inflammatory mediators were found to activate the HNSCC cell differentiation program.

**Conclusions:** Distinct cell differentiation states create a common tissue architecture in normal mucosa and HNSCCs. Our data demonstrate a loss of cell malignancy upon HNSCC cell differentiation, indicating that targeted differentiation approaches may be therapeutically valuable. Moreover, we describe KRT17 to be a candidate biomarker for HNSCC cell differentiation and early tumor detection.

## Introduction

Head and neck squamous cell carcinoma (HNSCC) is a highly malignant disease that represents the 6th most common type of cancer in the world. Five-year survival rates of about 50% have remained unchanged for decades [1, 2]. Despite recent advances in molecular targeting and immunotherapy, the prognosis of affected patients remains devastating. Even though new approaches in immunotherapy and molecular therapy exist, the standard of care for HNSCC remains surgery, cytotoxic chemotherapy, and radiotherapy. These treatments show limited tumor-specificity and thus frequently damage many vital structures in the head and neck area, leading to high morbidity even in long-term survivors [3]. Hence, new tumor-specific therapy approaches are urgently needed.

Squamous cell carcinomas are characterized by cornified areas detectable by histopathology. Cornification represents the normal differentiation path of keratinocytes in the epidermis and the oral mucosa and ensures the formation of a protective barrier on our body’s surface [4]. The process of differentiation begins with cells in the stratum spinosum exiting the cell cycle and strengthening their cytoskeleton. This leads to detachment from the basal membrane. In the next layer, the stratum granulosum, keratinocytes take on a flattened shape and increase the production of keratin and early cornification markers, such as loricrin and transglutaminase 3 (TGM3). These markers crosslink the cytoskeleton to form the cornified envelope. Additionally, lipid-filled cellular inclusions called laminar bodies form in the cytoplasm. Finally, the stratum corneum is formed, which displays late cornification markers like involucrin and SPRR3. Cells lose organelles and nuclei, becoming a dead shell with a lipid-encircled cornified envelope.[5]. In this way stem-like basal cells differentiate, renew the oral mucosa, and thereby create a protective barrier [4].

In the malignant context however, “cell differentiation” is defined by how closely a tumor cell resembles its original tissue, but little is known about the potential of long-term repopulating tumor cells to terminally differentiate and lose their malignancy due to epigenetic changes like chromatin remodeling processes. It is believed that the malignant cell state prevents terminal differentiation [6] or causes a plastic differentiation behavior with mixed or reversible differentiation phenotypes [7–10]. In this study, we have modeled HNSCC cell differentiation, describe differentiation markers, show close similarities in the tissue structure reflecting their normal tissuès counterpart, and established initial protocols to trigger this process. This may provide a rationale for the development of differentiation therapies driving HNSCC cells into cornification.

## Results

### Primary HNSCC spheroid models

The process of keratinization is a characteristic of squamous cell carcinomas. To study this differentiation path in HNSCC, we employed the primary spheroid model S18 derived from human papillomavirus (HPV)-negative oropharyngeal HNSCC of a previous study [11], further referred to as patient 1 (P1). Moreover, we established a primary cell culture from a xenograft tumor model of another study [12], derived from a second oropharyngeal HPV-negative HNSCC (P2; suppl. Table S1). The xenograft tumors of P2 closely resembled the original tumor in histology (suppl. Figure S1A) and adherent cells were isolated from the xenograft tumor tissue that could be expanded for more than 20 passages in stem cell medium without phenotypic changes (SCM, suppl. Figure S1B). The cells formed a spindle-like network of adherent cells that released spheroids into the supernatant (Figure 1A, suppl. Figure S1B). After spheroids were harvested from the supernatant, detached, and re-plated into normal flasks, cells regenerated the adherent spheroid-forming cell layer (suppl. Figure S1C), indicating that the spheroid cells as well can repopulate the original cell culture.

**Figure 1:**
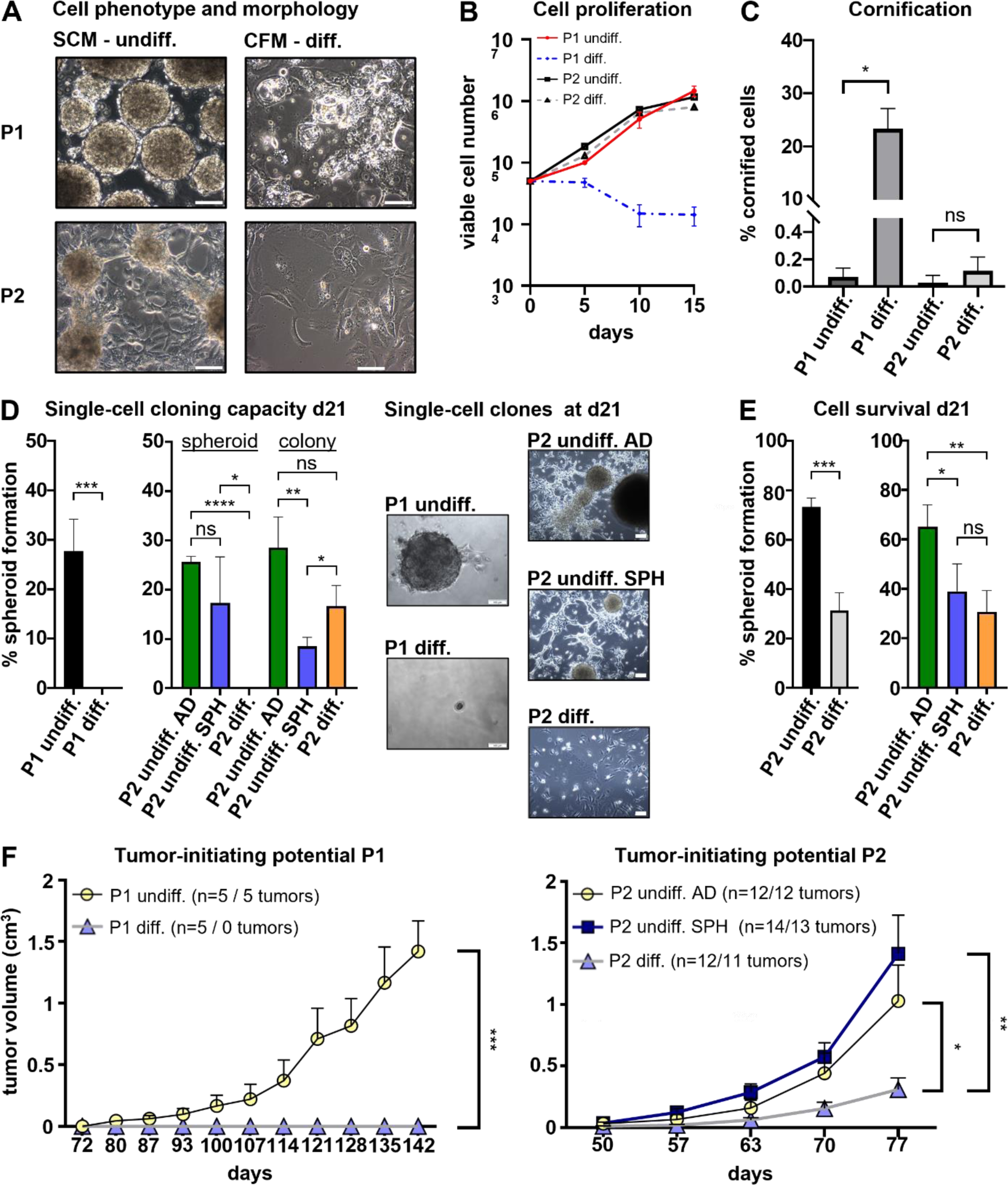
A HNSCC differentiation model reveals a loss of cell malignancy. **(A)** Cells of P1 grow as spheroids in SCM (undifferentiated), whereas P2 cultures display a loosely adherent cell layer that forms spheroids and releases them into the culture medium. Treatment of SCM cultures with differentiation medium (CFM) results in the formation of an adherent layer of enlarged cells with irregular shapes. P1 cells show in CFM show cell inclusions and bubble-like structures reminiscent of cornification. **(B)** CFM-treated cells of P1 lose proliferation, whereas cells of P2 continue to expand. **(C)** Cornification assay reveals cornified envelope formation in CFM-treated cells of P1 only; *p = 0.0005; ns = not significant. **(D)** Single-cell cloning assay for 21 days (d21) confirms efficient self-renewal and clonal re-population capacity of P1 spheroid cells which was totally abolished in CFM. Adherent and spheroid cells of P2 in SCM were equally able to form spheroids from single cells. Spheroid cells in SCM showed a reduced tendency to form adherent colonies from single cells. Single cells of P2 in CFM could not form spheroids; *p < 0.05; **p < 0.01; ***p < 0.005; ****p < 0.0001; ns = not significant. Examples of representative clonal cultures at day 21 (d21) are displayed. **(E)** Treatment with differentiation medium for 21 days reduces the survival of single cell clones of P1 and P2; *p < 0.05; **p < 0.01; ***p < 0.001; ns = not significant. **(F)** Subcutaneous transplantation of undifferentiated and differentiated populations into immunodeficient NSG mice showed diminished tumor-initiating cell potential of CFM-treated cells of P1 and P2. No significant differences were observed between adherent (AD) and spheroid cells (SPH) in SCM; *p < 0.05; **p < 0.005; ***p < 0.0005. p-values were determined by student’s t-test at the endpoint of the experiment.

### HNSCC differentiation by cornification abolishes cell malignancy

A described protocol to interfere with tumor stem cell self-renewal and to induce differentiation is to change from serum-free culture conditions to serum-containing medium [9, 10, 13]. Cardiac fibroblast medium (CFM), a serum-containing medium, displayed striking differentiating effect on HNSCC cells (Figure 1A). Upon treatment with this medium, P1 cells fully adhered to the culture dish displaying variable but strongly increased cell sizes with pleomorphic cell shapes (Figure 1A) and lost proliferation (Figure 1B). P2 xenograft tumor cells also adhered and acquired enlarged pleomorphic cell bodies (Figure 1A) but continued to proliferate (Figure 1B). CFM-cultured cells of P1 developed thickened cell envelopes and cytoplasmic inclusions reminiscent of lamellar bodies, indicative of cornification. Indeed, we detected about 200x more cornified envelopes in CFM-cultured P1 cells than in control cells (Figure 1C). This was insignificant in P2. SCM-cultured cells of P1 expanded from single cells to whole spheroids/colonies with an efficiency of about 30% which was totally abolished in CFM. P2 cells were able to efficiently repopulate in both media. However, cells in CFM did not form spheroids but grew as adherent colonies, whereas single-cell clones of adherent and spheroid SCM cells both reestablished the mixed adherent/spheroid culture (Figure 1D). Moreover, CFM treatment compromised the survival of P1 and P2 cells (Figure 1E). Transplantation of P1 and P2 cells into immunodeficient mice resulted in efficient xenograft tumor formation from SCM cells, whereas CFM populations displayed strongly reduced (P2) or even annihilated (P1) tumor-initiating cell (TIC) capacity (Figure 1F). Within the mixed adherent/spheroid SCM culture of P2 adherent and spheroid cells showed no significant differences. In P1, histopathology analysis also revealed cystic tissue architecture with large areas of necrosis that more closely resembled lymph node metastases than the original tumor (suppl. Figure 2A). For P2, a similar cystic tissue architecture of xenograft tumors was observed (suppl. Figure 2B). CFM and SCM tumors matched in histology with insignificantly different extents of tissue cornification (suppl. Figure 2C) indicating that remaining TIC after differentiation treatment retained their imprinted differentiation behavior. These results show that the differentiation of primary HNSCC cells causes a loss of malignant properties and a phenotype reflecting normal corneocytes.

### HNSCC cell keratinization is associated with chromatin reorganization and wound healing-associated signaling

We performed ATAC-seq and RNA-seq to dissect the changes underlying tumor cell differentiation (Figure 2A). ATAC-seq confirmed a reduction of open chromatin regions (OCRs) in the genome upon differentiation by 64.6% in P1 cells and by 59.08% in P2 cells (Figure 2B, suppl. Table S2), indicating closure of the genome reminiscent of stem cell differentiation in other systems [14–16]. Despite this net loss of chromatin accessibility, the proportion of open small promoters (<1kb) increased 1.75-fold in differentiated P1 cells and 2.05-fold in P2 cells. Larger promoter regions (>1kb) were proportionally not affected (Figure 2C, suppl. Table S3-S5). In both patients, ATAC-seq displayed that induction of tumor differentiation made OCRs accessible, correlating with epithelial cell differentiation, cell adhesion, and stress-associated pathways which included the regulation of response to wounding, programmed cell death, response to TGF-β, regulation of JNK cascade, and transmembrane receptor tyrosine kinase signaling (Figure 2D, suppl. Tables S6-S7). Transcription factor footprinting unraveled the mechanisms involved in P1 cell cornification and included TGF-β signaling via R-SMAD proteins SMAD2/3/4, activator protein-1 (AP-1) signaling via c-JUN, JAK/STAT3 signaling, interleukin 6 (IL6) and the pro-inflammatory transcription factor C/EBPβ (Figure 2E).

**Figure 2:**
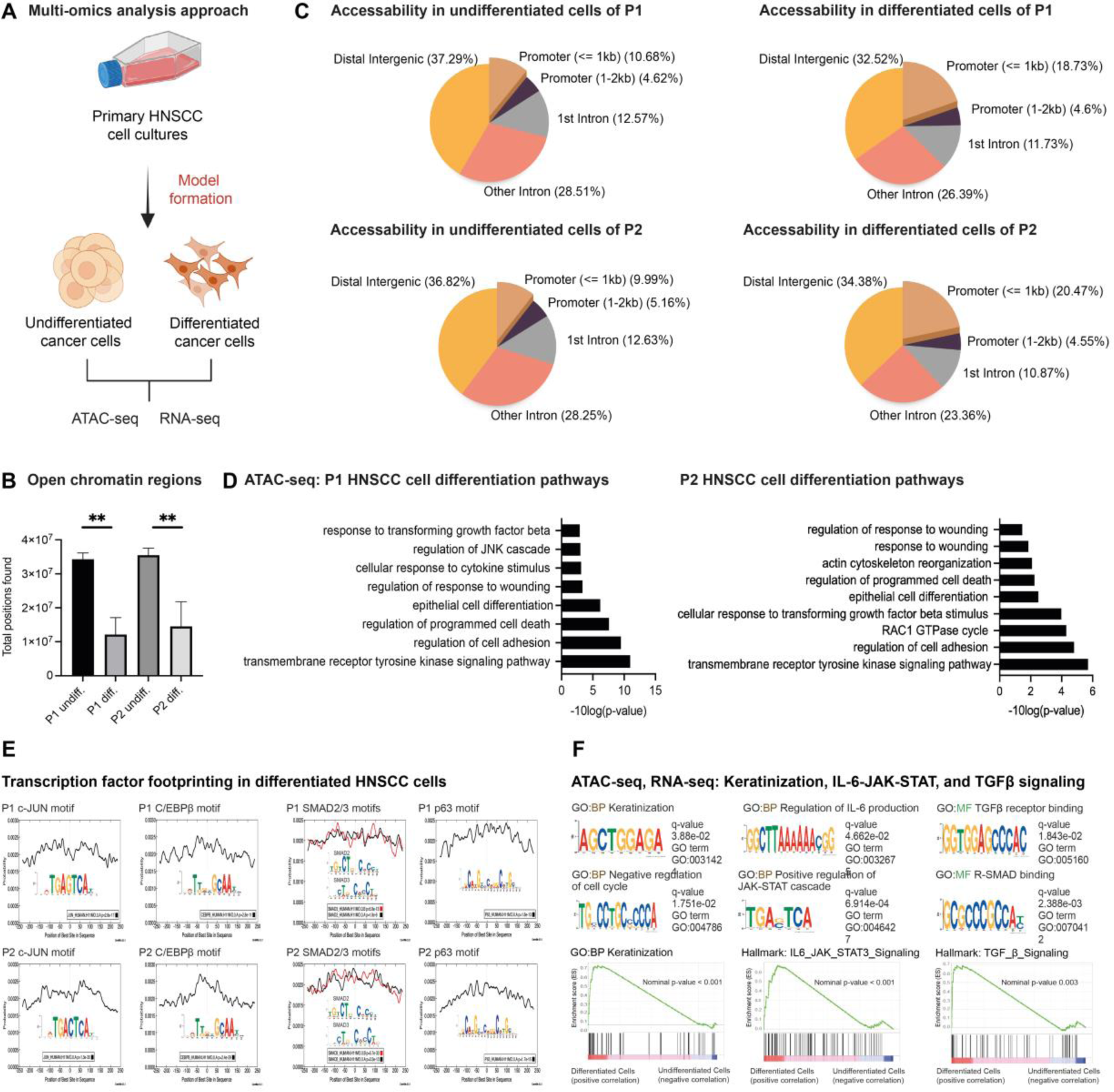
ATAC-seq identifies differentiation-associated signaling in HNSCC cells. **(A)** Overview of the omics-approaches to identify the changes underlying tumor cell cornification in HNSCC. **(B)** Genomic annotation of the OCRs in the cancer cells from both HNSCC patients showed a decrease in accessible areas upon differentiation (**p < 0.01) with technical replicates (n=3) and a statistical cut-off q-value of 0.05 for peak calling. **(C)** Despite an overall chromatin closure, the proportion of small open promoters (<1kb) increased. **(D)** GO terms display the relevant signaling pathways linked to the differential called OCRs within the differentiated cancer cells of both patients with Fisher’s exact test. **(E)** Transcription factor footprinting identified p63, c-JUN, C/EBPβ, and SMAD2/3 motifs in differentiating cancer cells. **(F)** Relevant pathways of HNSCC differentiation are Keratinization, IL6-JAK/STAT3, and TGF-β signaling, identified by gene ontology motif analysis (GOMo, ATAC-seq) and GSEA (RNA-seq); GO:BP = Gene Ontology Biological Process; GO:MF = Gene Ontology Molecular Function.

RNA-seq revealed hundreds of differentially expressed transcripts upon HNSCC cell differentiation (suppl. Tables S8-S11). Gene Set Enrichment Analysis (GSEA) confirmed the results of ATAC-seq identifying keratinization, IL6-JAK-STAT, and TGF-β to be involved in HNSCC cell differentiation (Figure 2F). In differentiated cells of P1, Gene Ontology (GO) analysis revealed upregulated pathways associated with the differentiation of keratinocytes and skin development (Figure 3A, suppl. Tables S12-S15). Simultaneously, cell cycle-related transcripts were downregulated consistent with the above-described loss of proliferation. In contrast, P2 exhibited variations in cell metabolism and biomolecule synthesis. Overall, cells of P1 and P2 showed little overlap in the transcriptional changes when put into differentiating conditions (Figure 3B), and epithelial differentiation was less intense in P2 compared to P1 cells (Figure 3C, suppl. Tables S16-S17). In comparison between P1 and P2, pathways related to keratinocyte differentiation, cytoskeleton organization, and keratinization were only upregulated in differentiating P1 cells, whereas cells of P2 up-regulated genes associated with cell migration, cell development, and regulation of signal transduction (Figure 3C).

**Figure 3:**
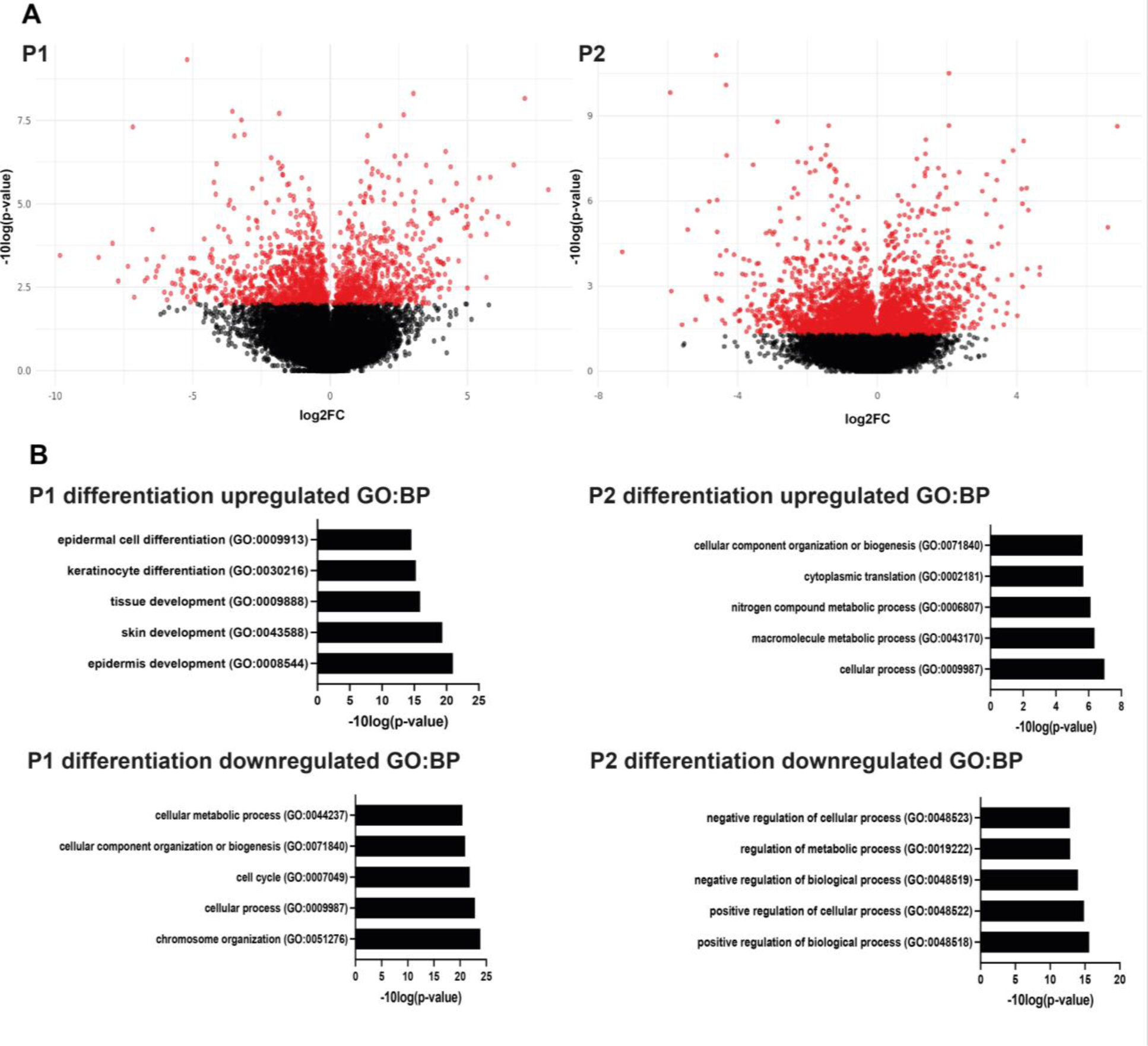
RNA-seq analysis reveals global expression changes upon HNSCC cell differentiation. **(A)** Volcano plot of global gene expression showing significantly up- and downregulated genes of differentiated and undifferentiated cells of two distinct HNSCC cases **(B)** Enrichment analysis shows the top five significant transcriptome changes with associated Gene Ontology Biological Process (GO:BP) pathways. Technical replicates were used (n=3, except P1 undifferentiated n=2) with a statistical cut-off p-value of 0.05.

As keratinization (or cornification) represented the main reaction of P1 cells to differentiating culture conditions, we searched for markers of this process. The cornification marker SPRR3 was 9.06-fold more accessible upon differentiation in P1 and upregulated 94.09-fold (suppl. Tables S4 and S8). KRT17 showed 14.59-fold increased accessibility and was upregulated 59.29-fold and displayed a striking accessibility peak only in differentiated P1 cells (suppl. Figure S3). In P2, KRT17 expression increased 2.8-fold (suppl. Table S12).

### High expression of KRT17 and SPRR3 marks HNSCC differentiation

As outlined above, ATAC-seq and RNA-seq analyses showed the upregulation of the basal cell marker KRT17 and small proline-rich proteins like SPRR3 in differentiating P1 cells which may serve as markers of HNSCC differentiation. IF analysis confirmed a strong upregulation of KRT17 and SPRR3 in CFM-treated P1 cells (Figure 4A). P1 spheroids stained predominantly negative for these markers. In P2, KRT17 was expressed by a subpopulation of cells under any condition, whereas SPRR3 was not detected (Figure 4B). To further investigate KRT17 and SPRR3 as markers of HNSCC differentiation, we stained the original tumor tissue of P1 and P2 and tissue of four additional patients with HPV-negative tumors of oropharyngeal and hypopharyngeal location (P3-6, suppl. Table S1). Normal mucosa attached to the surgically removed tumor piece showed KRT17 expression only in basal cells which was downregulated upon tissue maturation. SPRR3 initially displayed nuclear localization starting midway in the stratum spinosum followed by the formation of an SPRR3+ cell envelope towards the outer mucosal layers (Figure 4B-C). In contrast, KRT17 appeared to precede SPRR3 in the HNSCC tissue of all examined patients surrounding keratinized areas, as described previously [17], and to co-localize with SPRR3 towards the core of keratin pearls (Figure 4B-C). This indicates that KRT17 is a tumor-specific differentiation marker. P2 and P3 displayed hyperplastic/dysplastic tissue adjacent to the tumor. Interestingly, in both cases KRT17 expression pattern reflected the tumor and not the normal mucosa (Figure 4D) highlighting the reversal of KRT17 from a basal stem cell marker to cornification-associated differentiation-indicating protein to be an early event in HNSCC development. These data highlight KRT17 as a differentiation marker in HPV-negative HNSCCs and propose it as a possible biomarker for early tumor detection.

**Figure 4:**
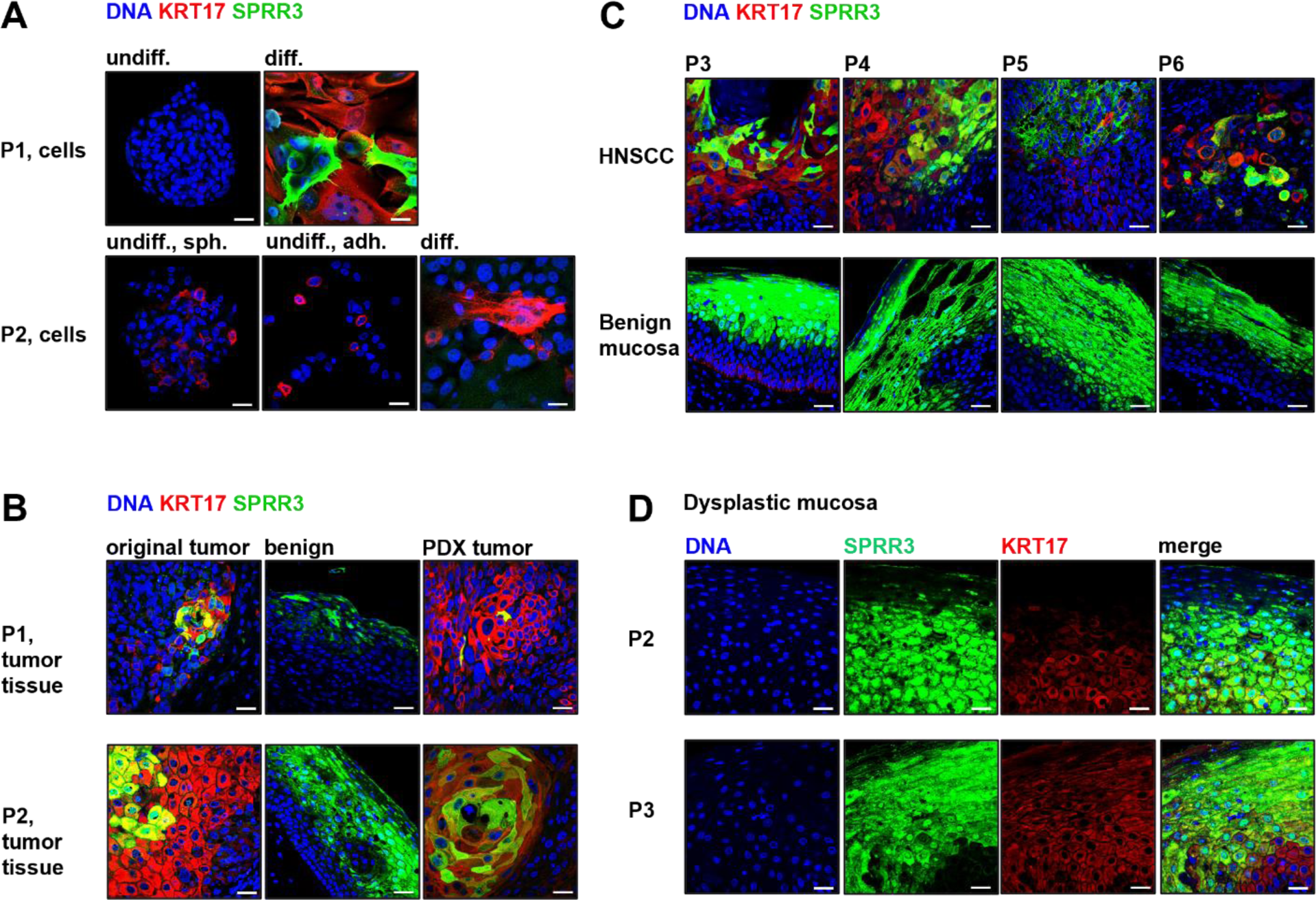
KRT17 expression indicates premature differentiation stages in HNSCC tissue and dysplastic mucosa. **(A)** P1 cells upregulate KRT17 and late cornification marker SPRR3 upon treatment with differentiation medium (CFM) while undifferentiated spheroid cells in stem cell medium (SCM) stain negative for this marker. In P2, KRT17 expression is restricted to a subpopulation in both media which was also observed in adherent (adh.) and spheroid (sph.) cells in SCM. **(B)** KRT17 was strongly expressed around SPRR3+ patches in the original patient’s HNSCC tissue and xenograft tumors in immunodeficient mice of P1 and P2, but not in normal mucosa of those patients. KRT17 seems to precede SPRR3 expression in human HNSCC tissue. **(C)** The tumor-specific association of KRT17 and cornification indicated by SPRR3 was reproduced in four additional patients (P3-P6). In benign mucosa of the same individuals KRT17 was barely visible in the basal layer only or not detectable, when imaged using the same instrument settings as used for the tumor sections. **(D)** Dysplastic mucosa attached to the tumor tissue of P2 and P3 showed a more abundant expression of KRT17 compared to normal mucosa spreading from the basal layer into the SPRR3+ zone; staining by indirect immunofluorescence (IF); scale bars = 10 µm.

### HNSCC differentiation can be triggered by inflammatory mediators

KRT17 is described to be upregulated in keratinocytes affected by psoriasis, a chronic inflammatory disease mechanistically resembling wound healing [18]. Interestingly, psoriasis is driven by the activation of a multitude of signaling pathways and transcription factors including STAT3, C/EBPβ, and AP-1 triggered by damage- or pathogen-associated molecules released from dying keratinocytes and invading inflammatory immune cells. Thus, we investigated whether there is a mechanistic connection between the signaling in psoriasis and HNSCC cornification. In histology, we noticed that keratinization of original tumor tissue of P1 was more intense in lymph node metastases which generally show a more (pseudo-)cystic growth pattern than primary tumors in HNSCC with cysts containing keratin and cellular debris in most cases [19]. Interestingly, P1 primary tumor tissue showed only small patches of keratinization, whereas lymph node metastasis tissue displayed larger differentiated areas predominantly surrounding necrotic areas, which was also observed in tumor tissue of other patients (Figure 5A). IF staining revealed the expression of cornification marker SPRR3 precisely at the interface between necrotic cysts and the vital tumor tissue (Figure 5B). Thus, we hypothesized that cornification might be triggered by inflammatory signaling from a necrotic possibly immune cell-infiltrated environment. In normal skin and psoriasis, cytokines influence the differentiation of keratinocytes [18, 20]. Thus, we treated P1 cells with described cytokines in SCM. Inflammatory cytokines induced a differentiation phenotype in HNSCC spheroid cells (Figure 5C). Most effectively, the combination of EGF, IL1α, IL1β, IL6, IL17A, IL22, TGFβ, OSM, and TNFα induced adhesion and upregulated expression of adhesion marker ITGB1 and differentiation markers KRT17 and SPRR3 (Figure 5C-D).These data show that differentiation can be imposed on HNSCC cells by the targeted application of inflammatory mediators.

**Figure 5:**
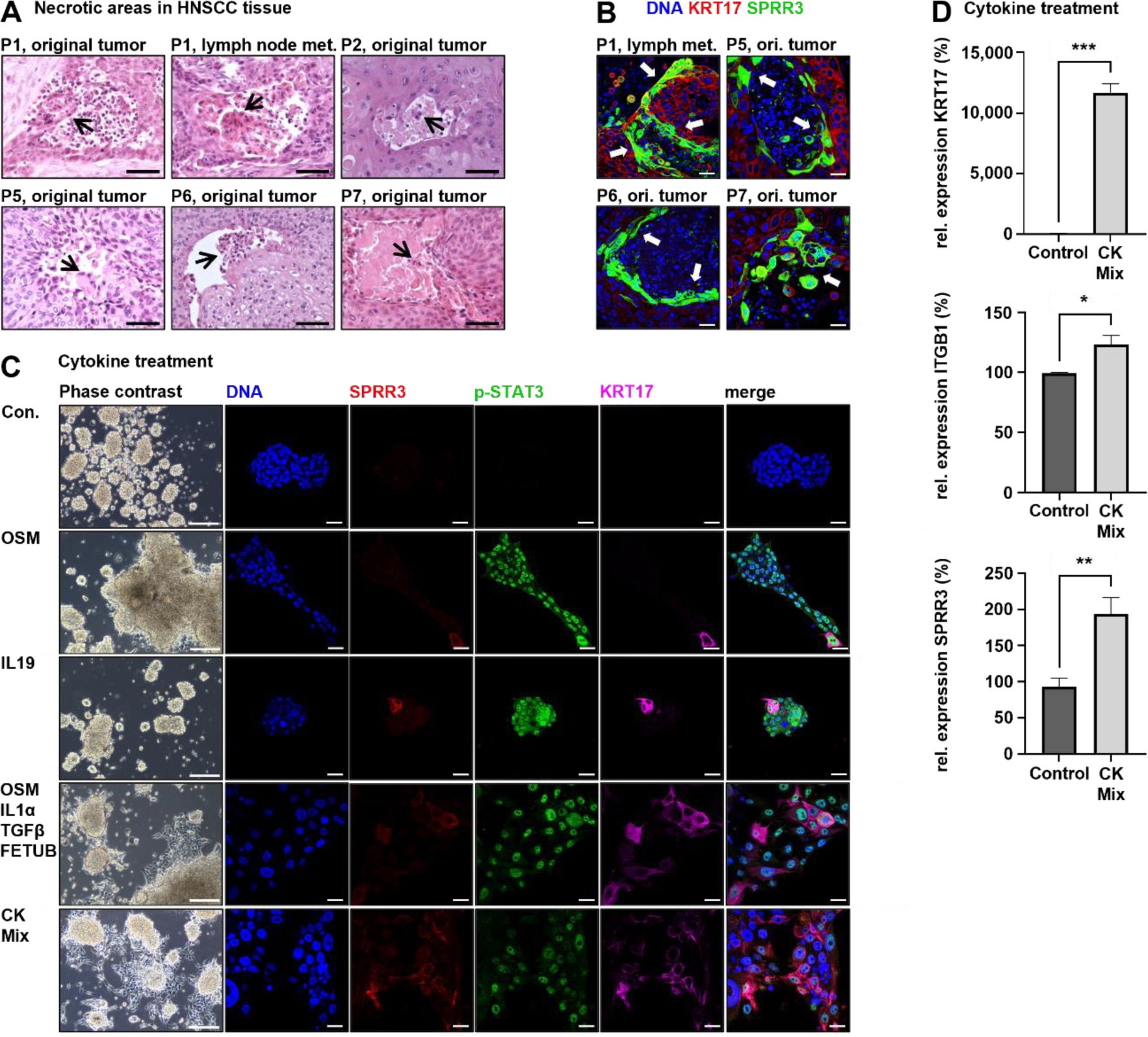
Inflammatory cytokines trigger differentiation in HNSCC cells. **(A)** HNSCC tissue shows cornified cells at the interface between vital tumor tissue and necrotic/immune cell-infiltrated areas and dead cornified cells in the necrotic cavity (arrows); scale bars = 50 µm. **(B)** IF analysis confirms the upregulation of cornification marker SPRR3 at the edge of necrotic/immune cell-infiltrated areas; scale bars = 10µm. **(C)** Treatment of undifferentiated HNSCC cells with combinations of inflammatory cytokines results in cell adhesion, activation of signaling through phosphor-STAT3-Y705 (pSTAT3), and upregulation of differentiation marker KRT17 and cornification marker SPRR3 compared to the untreated control (Con.). Single cytokines like IL19 and OSM were able to induce this differentiation phenotype, but higher numbers of cytokines in combination worked increasing effective; cytokine mix (CK Mix) = EGF, IL1α, IL1β, IL6, IL17A, IL22, TGFβ, OSM, and TNFα. Cells were treated for 5 days in SCM without hydrocortisone; cytokine concentration = 30 ng/ml; scale bars phase contrast = 200 µm; scale bars IF = 10 µm. **(D)** RT-qPCR shows increased relative expression of adhesion marker ITGB1, KRT17, and SPRR3 compared to the untreated control. Cells were treated with CK Mix for 3 days in SCM without hydrocortisone; cytokine concentration = 30 ng/ml; *p < 0.01; **p < 0.005; ***p < 0.0001, determined by student’s t-test.

### HNSCC tissue keratinization recapitulates normal mucosa architecture

Above, we describe an *in vitro* model of HNSCC cell differentiation and identified a signaling signature that can be induced by inflammatory cytokines. To verify our results in a larger set of patients, we examined the activity of these pathways utilizing indirect immunofluorescence in the original tumor tissue of P1, P2, and five additional patients (P3-P7) with HPV-negative HNSCCs originating from oropharyngeal, hypopharyngeal, and laryngeal location (suppl. Table S1, suppl. Figure S4), as well as the xenograft tumor tissue of P1 and P2. This set of HNSCC tissue samples contained cases with intense cornification associated with abundant keratin pearls (P2-P5, P7) and such with rather single-cell keratinization (P1, P6, suppl. Figures S2 and S4). IF staining for c-JUN, C/EBPβ, basal stem-cell marker p63 [21], and proliferation marker Ki67 in combination with the differentiation markers KRT17 and SPRR3 revealed four different zones of marker expression in normal mucosa adjacent to the tumor (Figure 6A-D): 1.) p63+/KRT17+ basal cells showed no activity of any signaling pathway. 2.) Cells in the stratum spinosum directly above the basal layer express Ki67 and p63 but lose KRT17 expression and activate c-JUN and C/EBPβ as evident by nuclear localization of these effectors. About midway in this zone of intense signaling cells start to express SPRR3. 3.) Towards the stratum granulosum SPRR3 changes to a cytoplasmic location as part of the cornified envelope. Cell signaling diminishes gradually as cells shut down viable processes during cornification. 4.) In the stratum corneum cells display pyknotic nuclei and stain negative for any marker, presumably due to epitope masking by transglutaminase activity. When we compared this normal mucosal structure to keratinized areas in HNSCC, we observed that these zones were reflected in the tumor tissue (Figure 6A-D). Tumors with intense cornification showed a zone with small cells at the stromal interface lacking expression of c-JUN and C/EBPβ except for Ki67 and p63. The expression of p63 and Ki67 correlated in some patients. In closer proximity to keratin pearls, cells become larger with more cytoplasm and nuclei appearing hollow and spotted reminiscent of heterochromatin formation. These cells stain positive for c-JUN and C/EBPβ but lose Ki67 indicating loss of proliferation (Figure 6A-C). KRT17 and SPRR3 are gradually upregulated in this zone. At the interface between this zone and destained fully keratinized tumor cells with pyknotic nuclei, cells reside displaying variable morphology and strong SPRR3 expression. Interestingly, in tissues with a low tendency to cornify (P1 and P6) these distinct differentiation states were also found but the spatial location of cells of each phenotype was rather chaotic. Unlike their organization into sequential layers in normal mucosa, in low-differentiated HNSCC tissue the distinct “layers” consisted of few or single cells only (Figure 6D). Altogether, HNSCC tissue mimics the differentiation architecture of its normal tissue counterpart (Figure 8).

**Figure 6:**
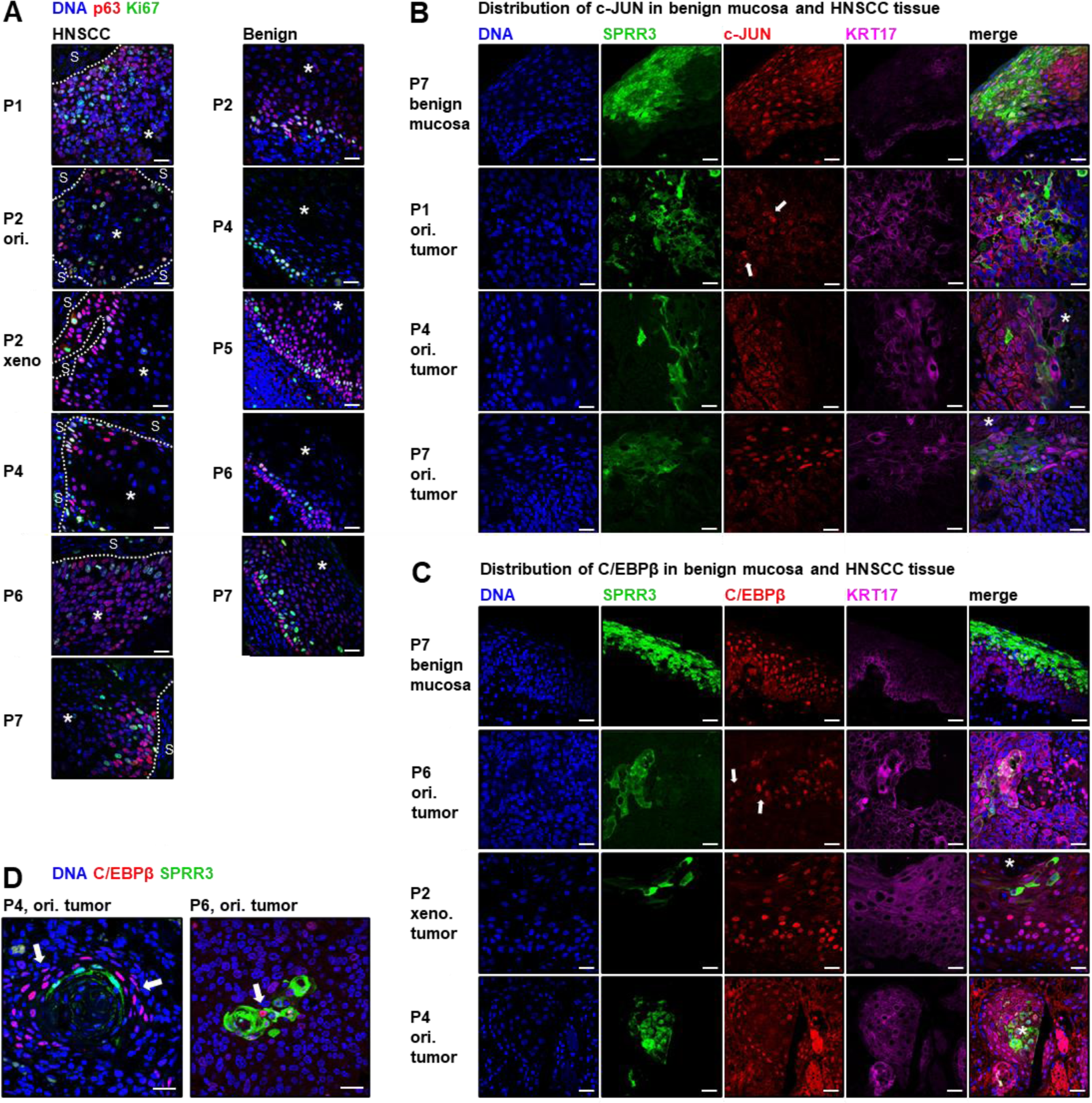
HNSCC tissue architecture resembles benign mucosa maturation. **(A)** Expression of basal keratinocyte marker p63 [21] and proliferation marker Ki67 were analyzed by indirect immunofluorescence. In benign mucosa, nuclear p63 is observed in the basal layer and in the lower stratum spinosum with Ki67+ cells located in a small layer above the basal cells. In HNSCC, a zone of cells positive for p63 and/or Ki67 is located at the tumor-stroma interface (dashed line, S = stroma) and both markers are downregulated towards the area of tumor differentiation (indicated by *). For example, P1 tumor tissue is largely undifferentiated, poorly organized, and shows abundant p63 and Ki67 expression which decreases surrounding the necrotic area. In contrast, HNSCCs of P2 and P4 show higher differentiation and p63/Ki67 expression is restricted to a small layer of cells. **(B-C)** HNSCC activates signaling via transcription factors c-JUN **(B)** and C/EBPβ **(C).** Representative examples are shown: each one benign mucosa sample, one poorly differentiated HNSCC (P1 or P6), and two higher differentiated HNSCCs (P2 and P4 or P4 and P7). Highly organized tumors and benign mucosa display a layer of cells with nuclear c-JUN and C/EBPβ right above the p63/Ki67+ layer detected in **(A)**. In these samples, basal cells in benign tissue and cells at the tumor-stroma interface in HNSCC show cytoplasmic location of these markers. Poorly organized tumors show fewer cells with nuclear c-JUN or C/EBPβ expression (arrows) which were mixed in between the SPRR3+ differentiated tumor cells indicating a perturbated layer organization. **(D)** Large (P4) and small (P6) areas of cornification harbor a zone of active signaling close to or within the SPRR3+ zone (indicated by nuclear C/EBPβ, arrows). Xeno. tumor = xenograft tumor tissue in immunodeficient mice; ori. tumor = original tumor tissue; dyspl. = dysplastic; scale bars = 10 µm.

## Discussion

### HNSCC cells recapitulate controlled stem cell differentiation

In this study, we modeled the transition of undifferentiated tumor-initiating cells to tumor corneocytes which was accompanied by an overall genome closure similar to stem cell differentiation in other systems [14–16]. Thus, the state of chromatin accessibility upon tumor differentiation was successfully altered, observing a significantly increased proportional accessibility of small promoter regions which may relate to the ongoing expression of housekeeping genes with simple promoters. This represents a remodeling process from euchromatin to heterochromatin, forming the differentiated state [22]. We conclude that HNSCC cell death by cornification is a controlled process similar to differentiation in normal tissue systems and that HNSCC cells can be forced to differentiate by epigenetic changes in a terminal way losing malignancy. To our knowledge, similar findings were only previously presented in neuroblastoma cells upon retinoid-induced ablation of their main oncogenic driver MYCN [23] but not in any type of carcinoma.

It is known that TIC formation and maintenance is controlled epigenetically [24] but little is described about which epigenetic mechanisms allow TICs or cancer stem cells to differentiate into bulk tumor cells. Single-cell RNA-seq revealed the differentiation dynamics of normal stem cells and confirmed that TICs display similar behavior upon transplantation in immunodeficient mice but in a skewed way frequently lacking large parts of normal lineage, displaying plasticity with less discrete differentiation stages, or acquiring irregular phenotypes [7]. Compared to single-cell techniques using human tissue, the bulk analysis of our model allows simultaneous analysis of the malignancy potential of distinct cell populations. In this way, we show a clear loss/reduction of clonal repopulation and tumor-initiating capacity upon forced tumor cell differentiation. This identifies the targeted induction of cornification as a promising future therapy approach.

### HNSCC differentiation activates wound-healing-like signaling

HNSCC cell differentiation *in vitro* directs activation of cell signaling via multiple pathways including but not limited to TGFβ, JNK/AP1, IL-6/JAK/STAT3, and C/EBPβ. We identify this signaling program to resemble wound healing and psoriasis [18]. Remarkably, these pathways are well-described as oncogenic in specific contexts, predominantly c-JUN [25], IL-6/JAK/STAT3 [26], and TGFβ [27]. C/EBPβ seems to exert an isoform-specific oncogenic role in breast cancer [28]. However, in certain malignancies, these molecules can exert anti-tumor properties in a context-dependent and tumor-stage-dependent manner as well. TGFβ is known to function as a tumor suppressor in early-stage tumor development and as an oncogene in later stages [27]. Other studies implicate c-JUN [29] and STAT3 [25] in tumor suppressive functions, too.

In HNSCC, increased nuclear pSTAT3 was shown to correlate with a dismal prognosis [30], and constitutive STAT3 activation is described as an early event in HNSCC development [31]. Moreover, high expression levels of c-JUN and p-c-JUN correlate with poor overall survival in hypopharyngeal squamous cell carcinoma [32]. Although these observations associate the expression of these factors with malignancy in HNSCCs, they do not prove their direct implication in cell self-renewal or any other malignant functions, as aggressive tumor stem-like cells may produce substantial amounts of differentiated bulk progeny.

### HNSCCs and benign mucosa share a common tissue architecture

Our data show that differentiation in HNSCC and normal mucosa is organized in layers of distinct cell populations. The activation of the wound-healing-like signaling discussed above appears to be the first step in cell differentiation in the benign and malignant context. The loss of Ki67 expression in this zone suggests a proliferation stop. In well-differentiated tumors, we see a basal-like aligned layer of Ki67+/p63+ cells located predominantly at the tumor-stroma interface. For P2 this was confirmed in the original tumor and the xenograft tumor. P1 displayed a rather undifferentiated tissue with most cells being p63+/Ki67+ indicating a high abundance of stem-like cells. P1 tissue rarely displays small patches of single-cell cornification with a wound healing-associated signaling zone consisting of individual cells surrounding SPRR3-positive cells. Thus, the differentiation structure of normal mucosa is retained in undifferentiated tumors but presents itself to be less obvious and chaotic. Remarkably, despite the low cornification in the original tumor tissue, the cultured cells of P1 were highly susceptible to differentiating stimuli. This indicates that certain undifferentiated HNSCCs may simply lack a differentiating microenvironment which could be altered therapeutically.

Interestingly, p63 is a known marker of basal stem cells in the normal skin and is essential for the proliferation and differentiation of keratinocytes [21]. It is known that the loss of p63 in normal skin changes the chromatin architecture and enhancer landscape [33–35] which correlates to the reorganization of the chromatin observed by us during HNSCC cell differentiation associated with decreasing nuclear p63 protein in differentiated areas of HNSCC tissue. Moreover, p63-dependent transcription of epidermal genes in keratinocyte differentiation depends on co-activator transcription factors and the activating epigenetic mark H3K27ac [33, 35, 36]. Thus, p63 is an important regulator of normal keratinocyte differentiation that according to our data might possess a similar role in HNSCC differentiation as well, which could be investigated in future studies.

### KRT17 is an early marker of HNSCC differentiation

Interestingly, RNA-seq analysis indicated KRT17 to be associated with HNSCC cell differentiation and we observed in various patients’ tissue that KRT17 changed from being a basal keratinocyte marker to an early differentiation marker preceding cornification in HNSCC. KRT17 is described as a stress-associated keratin in the skin and a marker of psoriasis, a chronic wound-healing disorder [18]. It is plausible that tumor cells upregulate stress-associated signaling as oncogenic genetic alterations cause abnormalities in cell metabolism, protein folding, and other vital processes. In tumor tissue however, KRT17 is well-known as an oncoprotein that promotes proliferation, immune evasion, survival, and other malignant hallmarks [37]. KRT17 confers increased resistance to cisplatin in cervical cancer [38] and bladder cancer [39] and elevated expression of this gene was described to correlate with poor survival in oropharyngeal squamous cell carcinoma [40]. These observations are in stark contrast to our results showing diminished tumor-initiating capacity of differentiated KRT17^high^ cells and an upregulation of this marker in proximity to cornified areas in the HNSCC tissue of distinct patients which is in line with other previous reports addressing the spatial expression pattern of KRT17 in HNSCC [17, 40]. Moreover, high KRT17 expression predicts better survival in HER2^high^ breast cancer [41] and is associated with lower invasion of tumor-promoting regulatory T-cells in HNSCC [42]. This indicates a context-dependent role of this marker in tumor biology. It is also possible that differentiated HNSCC cells unable to repopulate the tumor still facilitate oncopromoting functions like immunosuppression. Alternatively, tumors in advanced stages with higher tumor burden may contain more differentiated KRT17+ cells than smaller tumors due to more widespread necrosis, which we here link to higher differentiation in adjacent HNSCC tissue. This would associate malignancy to high KRT17 levels despite the lack of a causal connection. Moreover, our data show that dysplastic mucosa adjacent to the tumor reflects the tumor’s KRT17 expression pattern with a much stronger abundance of this marker, proposing KRT17 as a biomarker for early HNSCC detection.

### Differentiation therapies may be applicable in squamous cell carcinomas

In our study, we identify distinct HNSCC cell states within the differentiation process suggesting a rather hierarchical organization of long-term tumor repopulation. Tumor-initiating cell dynamics have been investigated previously in colon cancer (COCA) on a functional level. Using clonal marking experiments in serial transplantation in mice, researchers described a hierarchical organization of the colon tumor-initiating cell (TIC) compartment with a subpopulation of long-term repopulating cells and tumor-transiently amplifying cells (T-TACs) that lost self-renewal capacity and support tumor growth for a limited time frame [43]. This reflects the organization of normal colon regeneration with self-renewing LGR5+ stem cells at the bottom of the crypts and transiently amplifying cells to derive mature gut epithelial cells [44]. Moreover, another study revealed that colon cancer tissue is structured similarly to the normal colon with LGR5+ cells residing at the bottom of crypt-like patterns in the tumor tissue that give rise to KRT20+/LGR5-cells [45]. However, KRT20+ cells can efficiently repopulate the LGR5+ stem-like compartment upon depletion which demonstrates plasticity between the distinct COCA cell states and that differentiation in COCA is not terminal [8]. In another tumor type, pancreatic ductal adenocarcinoma (PDAC), long-term tissue repopulation was found to be similar in benign and malignant tissue with a clonal succession of transient TICs [9], reflecting the transient de-differentiation of acinar cells to regenerate defects in the normal exocrine pancreas [46]. Thus, COCA and PDAC TICs display plasticity in their phenotypic and functional differentiation which is believed to be a general phenomenon allowing tumors to avoid terminal differentiation by epigenetic mechanisms [24]. In contrast, our HNSCC differentiation model leads to cornification, which is terminal, and loss of malignant behavior, so differentiation therapies may be more efficient in this neoplasm than in other cancer types. Squamous cell carcinomas of different tissues of origin (e.g., lung or skin) may be vulnerable to such treatment as well.

Our data indicate that cytokines, possibly released by invading immune cells, might activate programmed cell death of tumor cells. Indeed, treatment with inflammatory cytokines induced adhesion and upregulation of differentiation markers in primary HNSCC cells. This suggests that activators of inflammatory signaling pathways and inhibitors of anti-inflammatory mechanisms in future may serve therapeutic purposes to trigger the differentiation of long-term repopulating cells in HNSCC.

## Materials and methods

### Human material and cell culture

Primary head and neck cancer tissue was obtained from medically indicated surgeries with informed consent of the patients, according to the declaration of Helsinki, and as approved by the ethics committee of the Ruhr-University Bochum (AZ 2018-397_3), as reported previously [11, 12]. The cells were cultured in PneumaCult™-Ex Plus Medium, referred to as stem cell medium (SCM, #05041, Stemcell Technologies, Vancouver, Canada) supplemented as described [11]. Adherent cells and spheroids were detached using Accutase (Capricorn Scientific, Ebsdorfergrund, Germany). Cells were differentiated in cardiac fibroblast medium (CFM, 316K-500, Cell Applications, San Diego, CA, United States) supplemented with 1% 200 mM L-Glutamine, 1% 100x Penicillin/Streptomycin, and 1% 250 µg/mL Amphotericin B solution (all three Capricorn Scientific). Dulbecco’s Modified Eagle Medium (DMEM, DMEM-HXA, Capricorn Scientific) was supplemented with 10% Fetal Bovine Serum Advanced (FBS Advanced, FBS-11A, Capricorn Scientific), L-Glutamine, Penicillin/Streptomycin, and Amphotericin B. Cells received fresh medium twice a week. Cells were analyzed for proliferation, and cornification, and treated with drugs and cytokines as outlined in supplementary methods.

### Mouse xenograft tumor models

All animal experiments were conducted as approved by the Institutional Animal Care and Use Committee of the University of Granada (procedures 12/07/2019/128) in accordance with the European Convention for the Protection of Vertebrate Animals used for Experimental and Other Scientific Purposes (CETS #123) and Spanish law (R.D. 53/2013).

NSG mice (NOD.Cg-Prkdcscid Il2rgtm1Wjl/SzJ, The Jackson Laboratory, Bar Harbor, ME, United States) were housed in appropriate sterile filter-capped cages with sterile food and water ad libitum. 1×10^6^ cells were transplanted subcutaneously as described previously [47], with differentiated and undifferentiated cell populations being injected side-by-side into the left and right flank of each mouse as performed in a previous study [9]. Animals were monitored for tumor development twice a week. Tumor sizes were measured using a vernier caliper and tumor volume was calculated by the formula (width×length2)/2. When the first tumor of each population reached the legal size limit, all mice of that experiment were measured and sacrificed. Tumors were harvested and fixed in 4% paraformaldehyde (PFA, 2.529.311.214, PanReac) for 24 hours. The fixed samples were embedded in paraffin following standard protocols.

### Histopathology analysis of HNSCC tissue

Paraffin-embedded tissue was sectioned to 2 µm thickness using a sliding HM430 microtome (Zeiss). Hematoxylin/Eosin (HE) staining was performed using standard protocols in a linear COT 20 tissue stainer (MEDITE, Burgdorf, Germany). HE stained sections were analyze by a senior pathologist (F.B.) to compare patient characteristics between original tumor and xenograft tumor models. To quantify keratinized areas in human tumor tissue every HE-stained section was subdivided into three random views and analyzed using an Axio Lab.1 microscope (Zeiss) and DISKUS software (Hilgers, Königswinter, Germany). Per view, the total area displaying malignant histology was determined in mm^2^. Next, the keratinized area within the tumor compartment was determined to calculate the proportional extent of cornification using Microsoft Excel.

### Indirect Immunofluorescence of HNSCC cells and tissue

Indirect Immunofluorescence (IF) of HNSCC cell cultures was performed as described previously [48]. Tumor tissue of patients and xenograft tumor models were analyzed by IF as established in prior studies [9]. Samples were imaged using an LSM780 confocal microscope (Zeiss, Oberkochen, Germany) and ZEN software (Zeiss). DNA was stained with DAPI (Sigma Aldrich, St. Louis, MO, United States) using 2µg/ml diluted in PBS + 0.1% bovine serum albumin (Capricorn Scientific). Utilized antibodies are listed in Supplementary Methods.

### RNA isolation and real-time quantitative PCR

Total RNA was isolated from cultured cells using innuPREP DNA/RNA Mini Kit (Analytik Jena, Jena, Germany) as recommended by the manufacturer’s protocol. RNA quality was determined with a BioDrop Duo+ spectral photometer (Biochrom, Holliston, USA). Real-time quantitative PCR (RT-qPCR) was conducted as described previously [11] using the following primers:

GAPDH-forward: CTGCACCACCAACTGCTTAG,
GAPDH-reverse: GTCTTCTGGGTGGCAGTGAT,
ITGB1-forward: GACGCCGCGCGGAAAAG,
ITGB1-reverse: TCTGGAGGGCAACCCTTCTT,
KRT17-forward: CCTCAATGACCAACACTGAGC,
KRT17-reverse: GTCTCAAACTTGGTGCGGAA,
SPRR3-forward: TGCACAGCAGGTCCAGCA,
SPRR3-reverse: GGCTCCTTGGTTGTGGGAA.

### Multi-omics analysis of HNSCC cell differentiation

Differentiated and undifferentiated HNSCC cell populations were examined by RNA-seq and ATAC-seq, as outlined in Supplementary Methods.

### Single-cell cloning assay

Cells were detached and seeded into U-bottom 96-wells at a density of approximately 1 cell/well in 100 µl volume of the desired medium. Plates were centrifuged and all wells were examined in a light microscope (Leica, Wetzlar, Germany) to identify wells containing single cells. These wells were examined after 21 days for survival and proliferation. Cells received fresh medium once a week. Obtained data were processed in Microsoft Excel (Microsoft Corporation, Redmond, WA, United States) and visualized using GraphPad Prism software (Graphpad Software Inc., San Diego, CA, United States).

## Acknowledgments

The authors thank Thorsten Seidel and the imaging facility of the Faculty for Biology of Bielefeld University for their support of this study.

The authors acknowledge the technical NGS team of the Omics CF NGS Unit and CeBiTec.

## Author contributions

Conceptualization, F.O.; methodology, F.O., S.G., L.M., K.N., J.K., D.L., and T.B.; validation, F.O., S.G., and L.M.; formal analysis, F.O., S.G., and L.M.; investigation, F.O., S.G., L.M., J.F., A.L., L.H., P.K., P.S., H.P., L.L., M.S., F.V.F., J.N., D.L., and F.B.; resources, L.U.S., F.B., G.E. K.N., J.K., T.B., and H.S.; data curation, F.O. and S.G.; software, S.G.; writing—original draft preparation, F.O. and S.G.; writing-review and editing, F.O., S.G., L.M., J.F., A.L., L.H., H.P., L.L., P.K., P.S., F.V.F., J.N., M.S., P.G., L.U.S., D.L., G.E., F.B., K.N., J.K., and T.B., and H.S.; visualization, F.O., S.G., and L.M.; supervision, F.O., S.G., G.E., T.B., K.N., J.K., and H.S.; project administration, F.O., L.U.S., G.E., F.B., T.B., K.N., J.K., and H.S.; funding acquisition, F.O., P.G., G.E. and H.S..

## Data and materials availability

All data not contained in this manuscript will be deposited in a publicly accessible database upon acceptance.

## List of Supplementary Materials

Supplementary Figure S1: HNSCC model of patient 2 (P2)

Supplementary Figure S2: Histology analysis of xenograft tumors in immunodeficient mice

Supplementary Figure S3: Accessibility of the KRT17 locus in undifferentiated and differentiated HNSCC cells.

Supplementary Figure S4: Histology of original tumor tissue of patients 3-7

Supplementary Figure S5: Architecture of oral mucosa and HNSCC tissue

Supplementary Table S1: Patient characteristics

Supplementary Table S2: Taq information of the differential peak analysis of the ATAC-seq

Supplementary Table S3: Genome peak distribution of the ATAC-seq from patient 1 and patient 2

Supplementary Table S4: The differential peak analysis of the ATAC-seq of patient 1

Supplementary Table S5: The differential peak analysis of the ATAC-seq of patient 2

Supplementary Table S6: ATAC-seq differential OCRs GO: BP patient 1

Supplementary Table S7: ATAC-seq differential OCRs GO:BP patient 2

Supplementary Table S8: The differential expression analysis of the RNA-seq of patient 1 diff. medium upregulated

Supplementary Table S9: The differential expression analysis of the RNA-seq of patient 1 diff. medium downregulated

Supplementary Table S10: The differential expression analysis of the RNA-seq of patient 2 diff. medium upregulated

Supplementary Table S11: The differential expression analysis of the RNA-seq of patient 2 diff. medium downregulated

Supplementary Table S12: RNA-seq diff. upregulated GO:BP patient 1

Supplementary Table S13: RNA-seq diff. downregulated GO:BP patient 1

Supplementary Table S14: RNA-seq diff. upregulated GO:BP patient 2

Supplementary Table S15: RNA-seq diff. downregulated GO:BP patient 2

Supplementary Table S16: The differential expression analysis of the RNA-seq of patient 2 versus patient 1 upregulated

Supplementary Table S17: The differential expression analysis of the RNA-seq of patient 2 versus patient 1 downregulated

Supplementary Table S18: Global proteome data from day 1 of patient 1 showing kinases only present in differentiated cancer cells

Supplementary Table S19: Global proteome data from day 4 of patient 1 showing kinases only present in differentiated cancer cells

Supplementary Table S20: Global proteome data from day 8 of patient 1 showing kinases only present in differentiated cancer cells

Supplementary Table S21: Global proteome data from day 1 of patient 2 showing kinases only present in differentiated cancer cells

Supplementary Table S22: Global proteome data from day 4 of patient 2 showing kinases only present in differentiated cancer cells

Supplementary Table S23: Global proteome data from day 8 of patient 2 showing kinases only present in differentiated cancer cells

Supplementary Table S24: Reactome of all kinases only present in differentiated cancer cells from patient 1

Supplementary Table S25: Reactome of all kinases only present in differentiated cancer cells from patient 2

Supplementary Methods

Supplementary References

