## Supplementary Material for "Inducible mucosa-like differentiation of head and neck cancer cells drives the epigenetically determined loss of cell malignancy"

**Supplementary Figure S1:** HNSCC model of patient 2 (P2)

**Supplementary Figure S2:** Histology analysis of xenograft tumors in immunodeficient mice

**Supplementary Figure S3:** Accessibility of the *KRT17* locus in undifferentiated and differentiated HNSCC cells.

**Supplementary Figure S4:** Histology of original tumor tissue of patients 3-7

**Supplementary Figure S5:** Architecture of oral mucosa and HNSCC tissue

**Supplementary Table S1:** Patient characteristics

**Supplementary Table S2:** Taq information of the differential peak analysis of the ATAC-seq

**Supplementary Table S3:** Genome peak distribution of the ATAC-seq from patient 1 and patient 2

**Supplementary Table S4:** The differential peak analysis of the ATAC-seq of patient 1

**Supplementary Table S5:** The differential peak analysis of the ATAC-seq of patient 2

**Supplementary Table S6:** ATAC-seq differential OCRs GO:BP patient 1

**Supplementary Table S7:** ATAC-seq differential OCRs GO:BP patient 2

**Supplementary Table S8:** The differential expression analysis of the RNA-seq of patient 1 diff. medium upregulated

**Supplementary Table S9:** The differential expression analysis of the RNA-seq of patient 1 diff. medium downregulated

**Supplementary Table S10:** The differential expression analysis of the RNA-seq of patient 2 diff. medium upregulated

**Supplementary Table S11:** The differential expression analysis of the RNA-seq of patient 2 diff. medium downregulated

**Supplementary Table S12:** RNA-seq diff. upregulated GO:BP patient 1

**Supplementary Table S13:** RNA-seq diff. downregulated GO:BP patient 1

**Supplementary Table S14:** RNA-seq diff. upregulated GO:BP patient 2

**Supplementary Table S15:** RNA-seq diff. downregulated GO:BP patient 2

**Supplementary Table S16:** The differential expression analysis of the RNA-seq of patient 2 versus patient 1 upregulated

**Supplementary Table S17:** The differential expression analysis of the RNA-seq of patient 2 versus patient 1 downregulated

**Supplementary Table S18:** Global proteome data from day 1 of patient 1 showing kinases only present in differentiated cancer cells

**Supplementary Table S19:** Global proteome data from day 4 of patient 1 showing kinases only present in differentiated cancer cells

**Supplementary Table S20:** Global proteome data from day 8 of patient 1 showing kinases only present in differentiated cancer cells

**Supplementary Table S21:** Global proteome data from day 1 of patient 2 showing kinases only present in differentiated cancer cells

**Supplementary Table S22:** Global proteome data from day 4 of patient 2 showing kinases only present in differentiated cancer cells

**Supplementary Table S23:** Global proteome data from day 8 of patient 2 showing kinases only present in differentiated cancer cells

**Supplementary Table S24:** Reactome of all kinases only present in differentiated cancer cells from patient 1

**Supplementary Table S25:** Reactome of all kinases only present in differentiated cancer cells from patient 2

**Supplementary Methods**

**Supplementary References**

**Please find Supplementary Tables S1-S25 in a separate document**

### Supplementary Figures

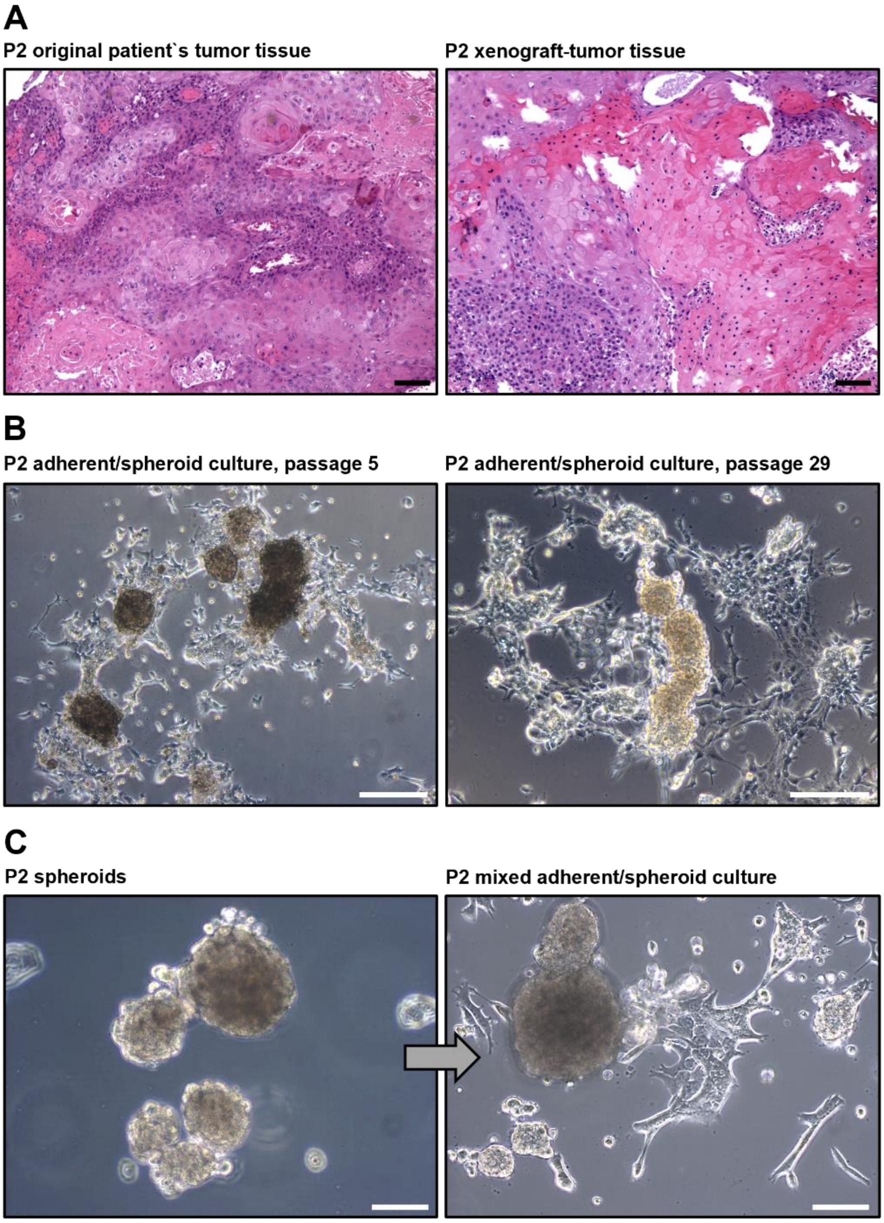


**Supplementary Figure S1: HNSCC model of patient 2 (P2). (A)** Original patient`s tumor and xenograft-tumor in immunodeficient NSG mice display similar histology; scale bars = 100 µm. **(B)** P2 cells from an adherent layer and give rise to spheroids that are gradually released into the culture medium; scale bars = 500 µm. **(C)** P2 spheroid cells can re-establish the mixed adherent/spheroid phenotype; scale bars = 100 µm.

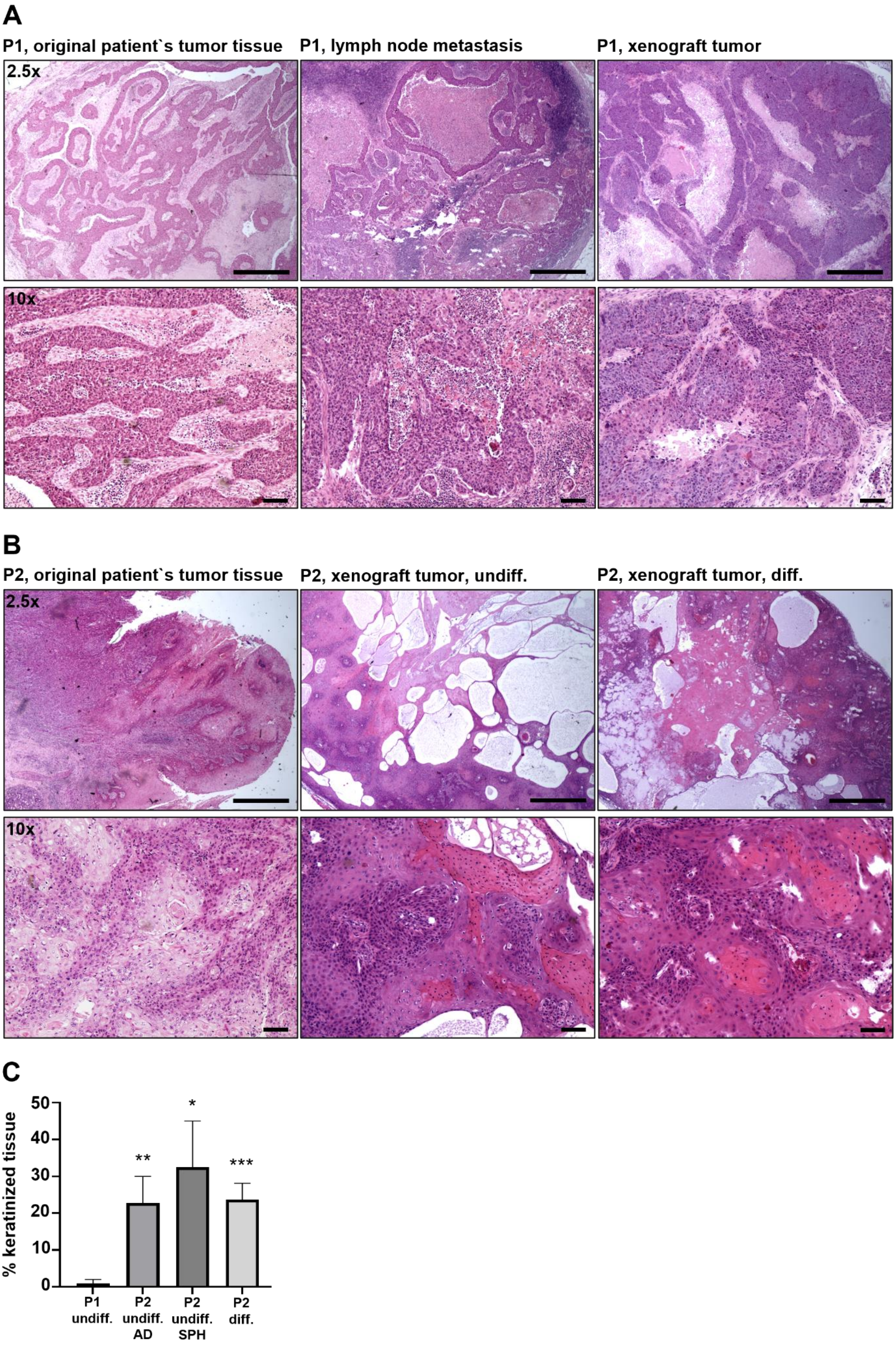
A
P1, original patient`s tumor tissue
P1, lymph node metastasis
P1, xenograft tumor
2.5x
10x
B
P2, original patient`s tumor tissue
P2, xenograft tumor, undiff.
P2, xenograft tumor, diff.
2.5x
10x
C
% keratinized tissue
50
40
30
20
10
0
**
*
***
P1 undiff.
P2 undiff. AD
P2 undiff. SPH
P2 diff.

**Supplementary Figure S2: Histology analysis of xenograft tumors in immunodeficient mice. (A)** HE-stained sections of P1 xenograft tumors display a cystic growth pattern with large areas of necrosis consistent with lymph node metastases of P1; scale bars = 1 mm upper panel, 100 µm lower panel. **(B)** P2 xenograft tumors show a cystic histology; scale bars = 1 mm. **(C)** Proportional quantification of keratinization in HNSCC xenograft tumor tissue of P1 and P2 in immunodeficient mice by histopathology analysis. All P2 populations were significantly higher keratinized than P1 tumors induced by SCM-cultured cells: *p < 0.01, **p < 0.005, ***p < 0.0005; determined by student`s t-test; n = 9 for P1 and n = 12 for each P2 population. Differentiation medium-treated (CFM) cells of P1 did not induce any xenograft tumors in mice in this experiment. All differences among P2 tumor populations were not significant. Significance levels were determined by student`s t-test; AD = adherent; SPH = spheroid.

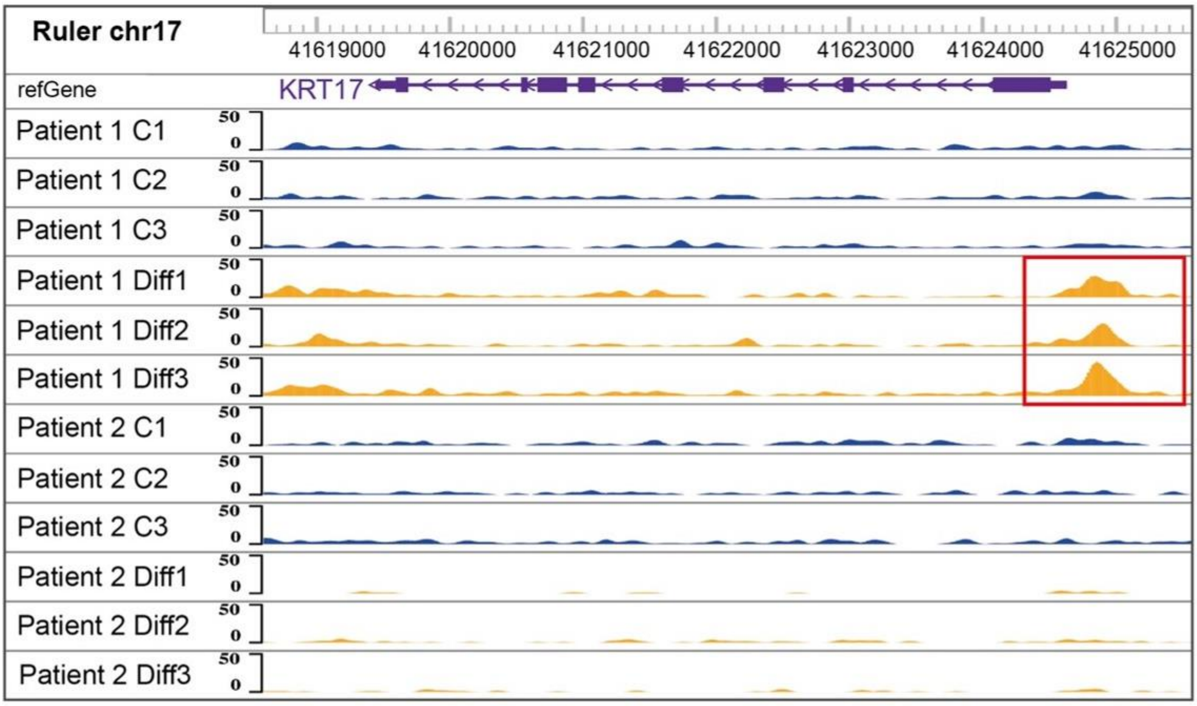


**Supplementary Figure S3: Accessibility of the *KRT17* locus in undifferentiated and differentiated HNSCC cells.** KRT17 was only accessible in highly sensitive cultures. Displayed are three independent samples of differentiated and undifferentiated cells of P1 and P2.

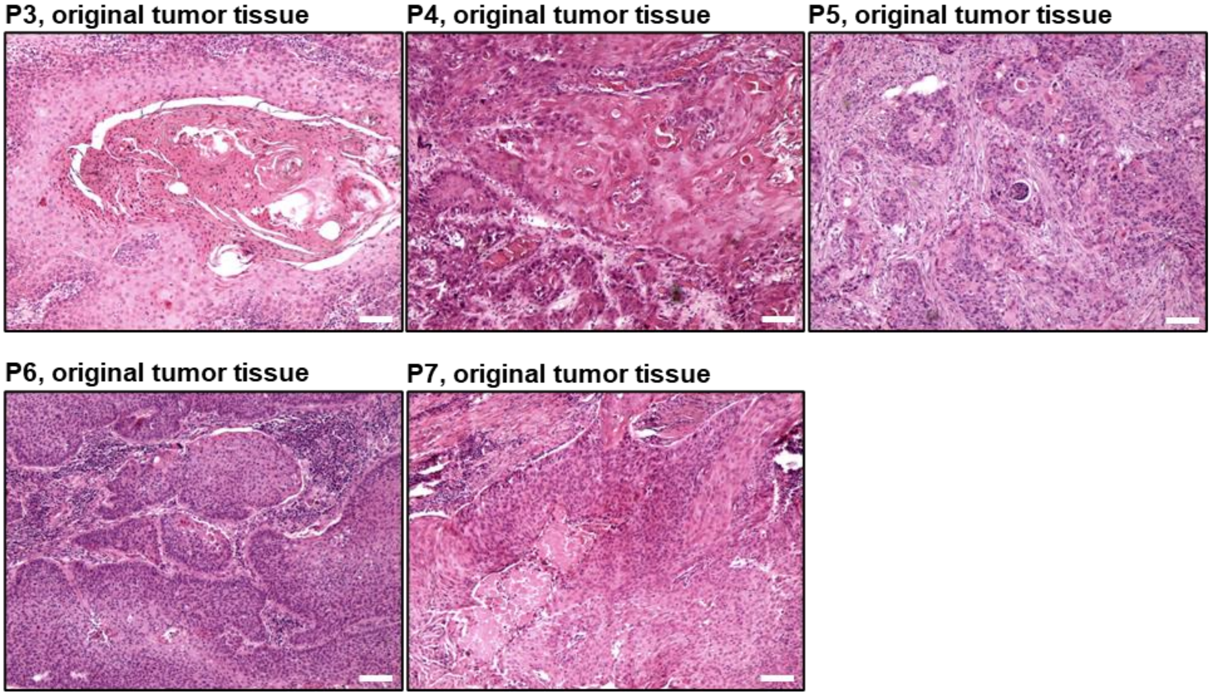


**Supplementary Figure S4: Histology of original tumor tissue of patients 3-7.** Hematoxylin and eosin staining; scale bars = 100 µm.

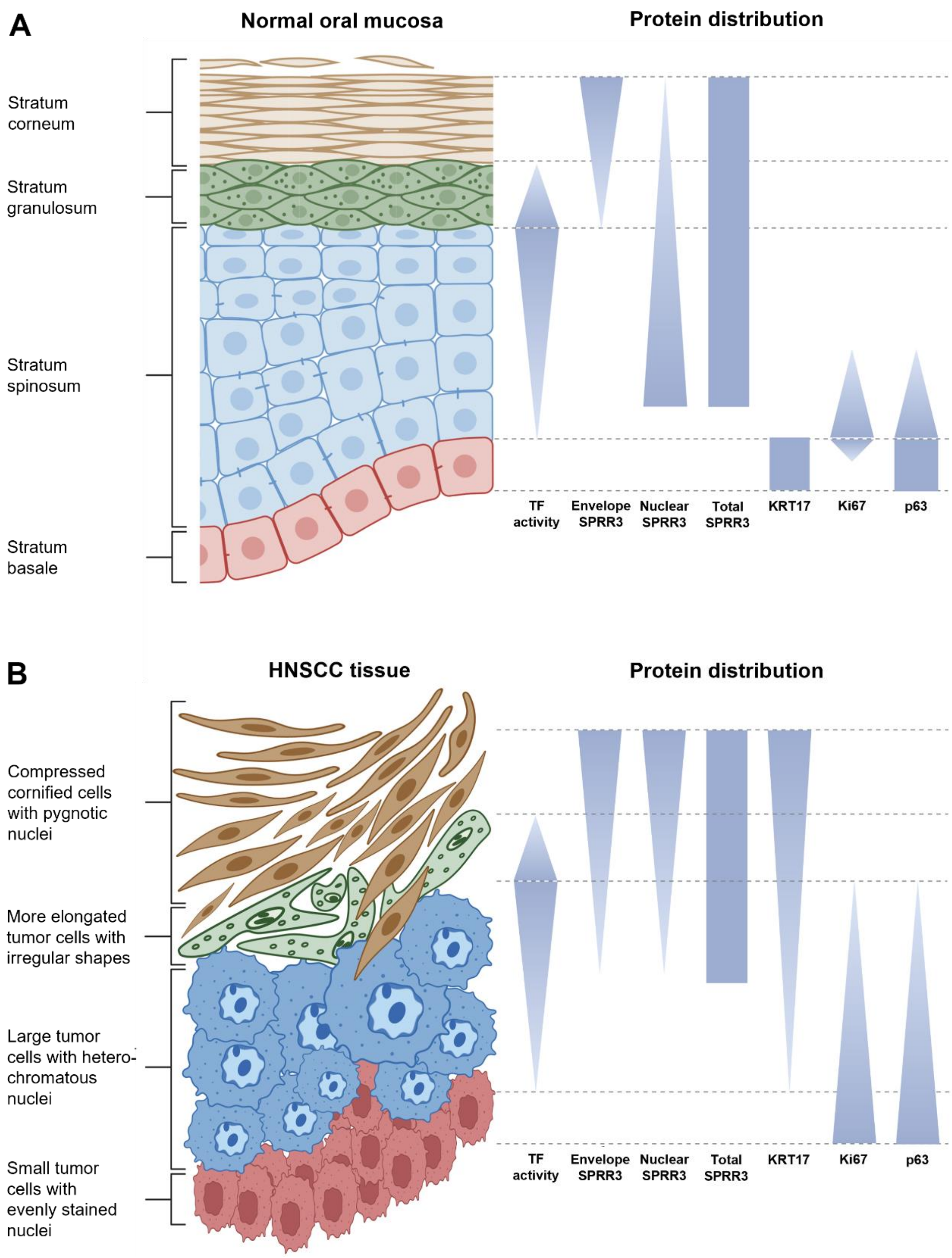


**Supplementary Figure S5: Architecture of oral mucosa and HNSCC tissue.** Schematic visualization of cell layer organization in **(A)** normal oral mucosa and **(B)** well-differentiated areas of HNSCC tissue. Protein marker distribution of KRT17, SPRR3, p63, proliferation marker Ki67 and transcription factors (TF) of the active signaling zone (e.g. C/EBPβ and c-JUN) is depicted. Please note that tumor tissue varies strongly in the proportional representation of each displayed layer. Created with Biorender.com.

### Supplementary Methods

#### Drug treatment of HNSCC cells

Small tumor spheroids (<100 µm diameter) were seeded in the appropriate medium in 12-well or 24-well cell culture dishes (STARLAB, Hamburg, Germany) containing 12 mm or 18 mm #1 cover slips (Carl Roth, Karlsruhe, Germany). Adherent cells were seeded at a density of 1-5x$10^4$ cells per well. Cells were treated with small molecules or recombinant human cytokines directly after seeding and incubated for 3 to 5 days. After that, cells were washed with phosphate buffered saline (PBS, Capricorn Scientific) and fixed with 4% PFA in PBS for 25 min. Cell populations were treated with EGF, IL1α, IL1β, IL6, IL17A, IL19, IL20, IL22, IL24, TGFβ, OSM, TNFα (all PeproTec, Hamburg, Germany), Fetuin-B (FETUB, NP_055190, R&D Systems/Bio-Techne, Abingdon, United Kingdom) or small molecules SB525334, IQ3 (both Selleck Chemicals, Houston, TX, United States), or C188-9 (MedChemExpress, Monmouth Junction, NJ, USA). Cytokine treatments were performed using SCM without hydrocortisone.

#### Antibodies used in indirect immunofluorescence analysis of cells and tissue

Primary antibodies: mouse-anti-human C/EBPβ (1:100, H-7, sc-7962), mouse-anti-human c-JUN (1:100, G-4, sc-74543), mouse-anti-human esophagin (SPRR3, 1:100, E-6, sc-514844), mouse-anti-human KRT17 (1:100, E-4, sc-393002), mouse-anti-human MEK-7 (MKK7, 1:100, E-7, sc-25288), mouse-anti-human p63 (1:100, D-9, sc-25268), mouse-anti-human SMAD2/3 (1:100, A-3, sc-398844, all Santa Cruz Biotechology, Dallas, TX, United States), rabbit-anti-human Ki67 (1:200, SP6, ab16667, Abcam), rabbit-anti-human SPRR3 (1:200, NBP2-13374, Novus

Biologicals/Bio-Techne, Abingdon, United Kingdom), rabbit-anti-human phospho-STAT3-Y705 (1:200, A16431, Antibodies Online, Aachen, Germany), rabbit-anti-human phospho-SMAD2-S465/S467 (1:50, E8F3R, 18338T, Cell Signaling Technologies), and guinea pig-anti-human KRT17; secondary antibodies: goat-anti-mouse-IgG-Alexa Fluor-555 (1:400, A21422), donkey-anti-rabbit-IgG-Alexa Fluor-488 (1:400, A11008), and goat-anti-guinea pig-IgG-Alexa Fluor-647 (1:400, A21450, all Thermo Fisher Scientific, Waltham, MA, United States).

**RNA Isolation**

Cells were cultured for eight days. Total RNA was isolated from cell culture (n=3) using innuPREP DNA/RNA Mini Kit (Analytik Jena, Jena, Germany) as recommended by the manufacturer's protocol. RNA quality was determined with a BioDrop Duo+ spectral photometer (Biochrom, Holliston, USA).

**RNA-seq Library Preparation and Sequencing**

PolyA-selected libraries were prepared from 200 ng of total RNA using QuantSeq 3′mRNA-Seq Library Prep Kit FWD for Illumina (Lexogen), according to the manufacturer's instructions. Size distribution and quality of the libraries were assessed by fragment analyzer (Agilent) and final libraries were sequenced in 75 bp single-end mode on a NextSeq2000 with P3 chemistry.

### Analysis of RNA-seq Data

RNA-seq analysis was performed with technical replicates (n=3, except BI18 undifferentiated n=2). The single-end reads were mapped to the human transcriptome (RefSeq Transcripts GRCh38; downloaded from NCBI) and quantified using Kallisto v0.44.0 [49]. Estimated counts were statistically analyzed using edgeR v3.28.1 [50]. The fold-change gene expression values were calculated in pairwise comparisons between the undifferentiated cancer cells as the control and differentiated cancer cells as treated. Genes were considered differentially expressed if their p-value was <0.05. Gene ontology analysis was processed using the GO web tool [51-53] with the human annotations applying Fisher's exact test. The GO annotation was visualized using GraphPad PRISM v9.0 (GraphPad Software, Inc). Gene set enrichment and leading-edge analyses were generated with GSEA software v4.2.1 [54, 55] with 1000 permutation numbers and a nominal p-value <0.05. The gene set enrichment map was generated using Cytoscape v3.9.1, with a cut-off value of p 0.01 [56].

### ATAC-seq Library Preparation and Sequencing

ATAC-seq analysis was performed with technical replicates (n=3). ATAC-seq was performed according to Buenrostro *et al.* [57] with harvested P1 and P2 cells, including isolation of intact nuclei and tagging with Tn5 transposase. After cell preparation, transposition reaction, library preparation, and purification, the quality of libraries was assessed using an Agilent Bioanalyzer High Sensitivity DNA Analysis kit. Sequencing was performed first on an Illumina MiSeq system in paired-end mode 2 x 50 bp and finally on the Illumina NextSeq 2000 system in paired-end mode 2 x 50 bp with P3 chemistry.

### Analysis of ATAC-seq Data

The ATAC-seq reads of each sample were mapped to the human reference genome GRCh38 using Bowtie2 v2.5.0 with default settings [58]. Samtools v1.16.1 was used for formatting, quality filtering, and removing duplicates [59] to guarantee unique mapping; the MAPQ (mapping quality, 0=non-unique, >10 probably unique) was set to a sophisticated value of 30. After examining each replicate separately, replicates of respected groups were combined to maximize the peak calling strength using the samtools merge function. MACS3 v3.0.0b1 was used for peak calling with the parameters "–nomodel –nolambda –keep-dup auto –call summits" the peaks were filtered by a q-value cutoff of 0.05 [60]. The sequences of peaks from 250 bp upstream and 250 bp downstream of the peaks were extracted with R v4.2.2 and Bedtools v2.30.0 [61] (R-Citation). In R v4.2.2 GenomicRanges v1.42.0 linked the ATAC-Seq peaks with the nearest genes. The promoter was defined as the region within 2 kb of the reference transcript start site [62]. Enriched motifs in the peak sequences were analyzed with MEME-ChIP v5.5.1 and default settings [63]. Gene ontology analysis of the identified motifs was done with GOMo v5.5.1 [64]. The differential peak analysis of the ATAC-seq was performed using HOMER2 v4.11 with a replicate FDR cutoff of 0.05 for peak identification calculated by DESeq2 v1.24.0 [65]. The genomic annotation of the peaks was determined using ChIPSeeker v1.34.1 [66, 67]. The genomic distribution of peaks was visualized using Adobe Illustrator 2023 (Adobe Inc., San Jose, USA).

### Cornification Assay

Cornification was quantified based on a previous protocol to isolate cornified envelopes [68]. Triplicates of P1 and P2 cells were seeded into 6-well plates in SCM or CFM until cells were 80-90% confluent. Cells were washed with 1x PBS, detached using Accutase (Capricorn Scientific, Ebsdorfergrund, Germany), and stained with trypan blue (Sigma Aldrich), followed by a count of live and dead cells using a Neubauer Chamber. The cells were washed in 1x PBS and subsequently mixed with 100 µL 4% SDS (Carl Roth GmbH, Karlsruhe, Germany) and 2% beta-Mercaptoethanol (Merck, Darmstadt, Germany) in PBS. The suspension is cooked at 95°C for 5-10 min. Lastly the cornified envelopes were counted using a Neubauer Chamber.

**Proliferation Assay**

Triplicates of P1 and P2 cells ($5x10^4$) were seeded into 12-well plate in SCM or CFM. After 5 days of incubation at 37°C with 5% $CO_2$ cells were washed with 1x PBS and detached using Accutase (Capricorn Scientific, Ebsdorfergrund, Germany). The cells were manually counted using a Neubauer Chamber. After that, cells were washed in 1x PBS, resuspended in culture medium and further incubated at the same conditions. The growth curves were generated over a period of 15 days, with repeated counting every 5 days.

51. Ashburner M, Ball CA, Blake JA, Botstein D, Butler H, Cherry JM, Davis AP, Dolinski K, Dwight SS, Eppig JT, et al: **Gene ontology: tool for the**

**unification of biology. The Gene Ontology Consortium.** *Nat Genet* 2000, **25:**25-29.

52. Gene Ontology C: **The Gene Ontology resource: enriching a GOld mine.** *Nucleic Acids Res* 2021, **49:**D325-D334.
53. Mi H, Muruganujan A, Ebert D, Huang X, Thomas PD: **PANTHER version 14: more genomes, a new PANTHER GO-slim and improvements in enrichment analysis tools.** *Nucleic Acids Res* 2019, **47:**D419-D426.
54. Subramanian A, Tamayo P, Mootha VK, Mukherjee S, Ebert BL, Gillette MA, Paulovich A, Pomeroy SL, Golub TR, Lander ES, Mesirov JP: **Gene set enrichment analysis: a knowledge-based approach for interpreting genome-wide expression profiles.** *Proc Natl Acad Sci U S A* 2005, **102:**15545-15550.
55. Mootha VK, Lindgren CM, Eriksson KF, Subramanian A, Sihag S, Lehar J, Puigserver P, Carlsson E, Ridderstrale M, Laurila E, et al: **PGC-1alpha-responsive genes involved in oxidative phosphorylation are coordinately downregulated in human diabetes.** *Nat Genet* 2003, **34:**267-273.
56. Shannon P, Markiel A, Ozier O, Baliga NS, Wang JT, Ramage D, Amin N, Schwikowski B, Ideker T: **Cytoscape: a software environment for integrated models of biomolecular interaction networks.** *Genome Res* 2003, **13:**2498-2504.
57. Buenrostro JD, Wu B, Chang HY, Greenleaf WJ: **ATAC-seq: A Method for Assaying Chromatin Accessibility Genome-Wide.** *Curr Protoc Mol Biol* 2015, **109:**21 29 21-21 29 29.
58. Langmead B, Salzberg SL: **Fast gapped-read alignment with Bowtie 2.** *Nat Methods* 2012, **9:**357-359.
59. Li H, Handsaker B, Wysoker A, Fennell T, Ruan J, Homer N, Marth G, Abecasis G, Durbin R, Genome Project Data Processing S: **The Sequence Alignment/Map format and SAMtools.** *Bioinformatics* 2009, **25:**2078-2079.
60. Zhang Y, Liu T, Meyer CA, Eeckhoute J, Johnson DS, Bernstein BE, Nusbaum C, Myers RM, Brown M, Li W, Liu XS: **Model-based analysis of ChIP-Seq (MACS).** *Genome Biol* 2008, **9:**R137.
61. Quinlan AR, Hall IM: **BEDTools: a flexible suite of utilities for comparing genomic features.** *Bioinformatics* 2010, **26:**841-842.
62. Lawrence M, Huber W, Pages H, Aboyoun P, Carlson M, Gentleman R, Morgan MT, Carey VJ: **Software for computing and annotating genomic ranges.** *PLoS Comput Biol* 2013, **9:**e1003118.
63. Bailey TL, Johnson J, Grant CE, Noble WS: **The MEME Suite.** *Nucleic Acids Res* 2015, **43:**W39-49.
64. Buske FA, Boden M, Bauer DC, Bailey TL: **Assigning roles to DNA regulatory motifs using comparative genomics.** *Bioinformatics* 2010, **26:**860-866.
65. Heinz S, Benner C, Spann N, Bertolino E, Lin YC, Laslo P, Cheng JX, Murre C, Singh H, Glass CK: **Simple combinations of lineage-determining transcription factors prime cis-regulatory elements required for macrophage and B cell identities.** *Mol Cell* 2010, **38:**576-589.

66. Yu G, Wang LG, He QY: **ChIPseeker: an R/Bioconductor package for ChIP peak annotation, comparison and visualization.** *Bioinformatics* 2015, **31:**2382-2383.

67. Wang Q, Li M, Wu T, Zhan L, Li L, Chen M, Xie W, Xie Z, Hu E, Xu S, Yu G: **Exploring Epigenomic Datasets by ChIPseeker.** *Curr Protoc* 2022, **2:**e585.

68. Jarnik M, Simon MN, Steven AC: **Cornified cell envelope assembly: a model based on electron microscopic determinations of thickness and projected density.** *J Cell Sci* 1998, **111 ( Pt 8):**1051-1060.
